# A new method to quantify the spatiotemporal localization of SnRK1.1

**DOI:** 10.1101/2025.04.14.648863

**Authors:** Candela Brugnara, María Candelaria Díaz, Julián Bultri, Daniela Liebsch, Francisco J. Hita, Dianela Aguilar Lucero, Corina M. Fusari, Jörn Dengjel, Valeria Levi, Nicolás E. Blanco

## Abstract

Maintaining energy homeostasis is a major challenge for plants in the current context of climate change. The Sucrose-non fermenting 1 (SNF1)-related kinase 1 (SnRK1) complex, a member of the SNF1-AMP-activated protein kinase (AMPK)-SnRK1 family of kinase complexes, is a central player in the regulation of cell energy homeostasis. The α-subunit of the complex, which possesses kinase activity and is known as SnRK1.1 or KIN10, plays a role in sensing energy status and coordinating metabolic reprogramming to counter any energy imbalance. The discovery of a dual and dynamic intracellular distribution of SnRK1.1 suggests that the activity and function of SnRK1 might be regulated by spatiotemporal changes.

To investigate the spatiotemporal distribution of SnRK1.1, we developed a protocol to quantify its intracellular distribution using fluorescence confocal images acquired along the z-axis in plants expressing SnRK1.1–eGFP. Using the open-source software Fiji/ImageJ, we calculated the ratio between nuclear and non-nuclear SnRK1.1 fractions and defined this as the N/ER index. We validated our method by analyzing the response of SnRK1.1 to photosynthesis inhibition by DCMU, including changes in protein levels and phosphorylation status. In addition, comparison with results obtained using a commercial software-based approach confirmed the compatibility of the N/ER index with different segmentation and quantification tools. Originally designed for leaf tissue images, this protocol can be broadly applied to assess the role of intracellular spatiotemporal changes in a wide range of kinases or fluorescently tagged recombinant proteins. Finally, SnRK1.1 intracellular distribution may also serve as a proxy to assess changes in cellular energy status.

**One sentence summary:** New method to track SnRK1.1 distribution and changes in plant cell energy status

## Introduction

Changes in growing conditions profoundly affect the growth and development of plants. The increasing frequency of unexpected climatic events negatively impacts on central plant processes like photosynthesis, ultimately affecting growth and development (Becklin *et al*., 2021). To sustain energy homeostasis and buffer energy imbalances caused by non-optimal conditions, plants rely on several intracellular mechanisms (Xiao *et al*., 2024). Among these, the sucrose non-fermentable 1 (SNF1)-related protein kinase 1 (SnRK1) complex plays a central role (Wurzinger *et al*., 2018; Peixoto and Baena-Gonzalez 2022). SnRK1 shares commonalities with its orthologs in yeast (SNF1) and in mammalian cells, the AMP-dependent protein kinase (AMPK) (Broeckx *et al*., 2016; Emanuelle *et al*., 2016). In response to decreases in cellular energy levels, SNF1/SnRK1/AMP kinase complex family orchestrates a shift toward catabolic reactions while suppressing energy-consuming anabolic processes, generating an energy surplus to withstand adverse growth conditions (Broeckx *et al*., 2016). However, differences between the complexes of photosynthetic and non-photosynthetic organisms suggest plant-specific adaptations in bioenergetic regulation. For instance, SnRK1 is a heterotrimeric kinase complex that contains a βγ scaffold subunit instead of the canonical γ subunit found in SNF1 and AMPK complexes (Emanuelle *et al*., 2015; Emanuelle *et al*., 2016). The presence of the plant-specific βγ subunit in the plant holoenzyme promotes a distinct folding and regulation of the other α and β subunits. In the case of the catalytic α subunit, the spatial organization of its “activation loop” (or “T-loop”), a domain whose phosphorylation promotes kinase activity, makes it less accessible and more resistant to dephosphorylation than in non-photosynthetic counterparts. (Emanuelle *et al*., 2015). Additionally, the T-loop in SnRK1.1 lacks the regulatory conformational changes observed in AMPK and SNF1 upon binding of AMP (Shen *et al*., 2009; Gowans *et al*., 2013), or AMP and ADP (Xiao *et al*., 2011), respectively (Oakhill *et al*., 2012; Emanuelle *et al*., 2015; Emanuelle *et al*., 2016). Therefore, SnRK1.1 kinase is likely to be constitutively active, as its T-loop remains predominantly phosphorylated (Ramon *et al*., 2019). Together, these peculiarities of the SnRK1.1 kinase suggest a unique regulatory mechanisms for its activity complex reflecting the plant-specific lifestyle (Roustan *et al*., 2016; Crepin and Rolland 2019; Peixoto and Baena-Gonzalez 2022).

The discovery of different intracellular SnRK1.1 fractions in plant cells has been crucial for studying the SnRK1-mediated signalling pathway from a spatiotemporal perspective (Gutierrez-Beltran and Crespo 2022). SnRK1 has been defined as a sensor or integrator of stress signals, and as a coordinator of the response, the latter function being related to the nuclear regulation of transcription (Baena-Gonzalez *et al*., 2007). The nuclear fraction of SnRK1.1 has been linked to the phosphorylation and regulation of the activity of transcription factors and the concomitant regulation of gene expression (Ng *et al*., 2013; Mair *et al*., 2015; Muralidhara *et al*., 2021; Peixoto *et al*., 2021; Henninger *et al*., 2022).The identification of a second fraction of SnRK1.1 associated with the ER indicates a separation of function among different intracellular SnRK1.1 pools, suggesting that the ER might be a point of gauging and/or integration of low-energy signals (Jamsheer *et al*., 2018b; Blanco *et al*., 2019; Crepin and Rolland 2019; Ramon *et al*., 2019; Gutierrez-Beltran and Crespo 2022). In favour of this hypothesis, the ER has been identified as the colocalization site of SnRK1.1 with some of its interactors, including DUF581-FLZ family proteins (Jamsheer *et al*., 2018a; Jamsheer *et al*., 2022), class II T6P synthase (TPS)-like proteins (Van Leene *et al*., 2022) and members of the TOR complex (Nukarinen *et al*., 2016). Furthermore, in leaves sections, changes in the SnRK1.1 fraction at the ER were triggered by different blockage of photosynthetic electron flow, which affects energy status (Blanco *et al*., 2019). Depending on the final redox status of the electron transport chain, either overoxidized by DCMU or overreduced by DBMIB, SnRK1.1 fraction either delocalizes from the ER or rearranges into bright cytosolic puncta. These studies plus the changes in the SnRK1.1-eGFP localization in mesophyll protoplasts expressing different β subunits of SnRK1 complex (Ramon *et al*., 2019) and in root cells in response to ABA treatment (Belda-Palazon *et al*., 2022) indicate a likely specific role of the different SnRK1.1 fractions (Blanco *et al*., 2019; Gutierrez-Beltran and Crespo 2022). Although the underlying mechanism has not been fully elucidated, growing evidence supports a role for the ER-associated SnRK1.1 fraction as a sensor of energy fluctuations in a standby state. This suggests that SnRK1.1 activation is spatially and temporally regulated, an essential factor that must be considered when investigating mechanisms of energy homeostasis.

The participation and activity of SnRK1 in cellular processes is currently determined by two methodological approaches based either on the analysis of the target genes or on the phosphorylation status of target proteins (Baena-Gonzalez *et al*., 2007; Peixoto *et al*., 2021; Van Leene *et al*., 2022). In the first case, the quantification of the expression levels of SnRK1.1 target genes (e.g., *SENESCENCE-ASSOCIATED PROTEIN 5 (SEN5), DARK INDUCIBLE 1* (*DIN1/SEN1*) and *6/ASPARAGINE SYNTHASE 1 (DIN6/ASN1)*, and *PROLINE DEHYDROGENASE 1* (*PRODH*)) have been used as proxies of the SnRK1.1 activity (Baena-Gonzalez *et al*., 2007; Pedrotti *et al*., 2018; Peixoto *et al*., 2021; Peixoto and Baena-Gonzalez 2022). Regarding the SnRK1 kinase activity, the identification of the SnRK1.1-dependent phosphoproteome or protein-protein interactions studies complemented by detection of target phosphopeptides have been used to verify SnRK1 activity (Cho *et al*., 2016; Nukarinen *et al*., 2016; Carianopol *et al*., 2020; Van Leene *et al*., 2022). Additionally, two methods to assess SnRK1 activity have also exploited its kinase activity *in planta* (Muralidhara *et al*., 2021; Sanagi *et al*., 2021; Avidan *et al*., 2023; Safi *et al*., 2023). Sanagi and co-workers generated Arabidopsis lines expressing a synthetic peptide derived from the rat Ser79 phosphorylation site of ACETYL COA CARBOXYLASE 1 (ACC), a conserved direct phosphorylation target of AMPK/SNF1/SnRK1.1 (Sanagi *et al*., 2021). Relative SnRK1 activity is determined by the quantification of immunoblot signals detected by antibodies anti-ACC pS79 compared to those of anti-HA or GFP, both domains included in the reporter construct. Different modifications of the technique have been used to determine the relative activity of the SnRK1 nuclear fraction by adding a localization sequence (NLS) to the AAC-peptide (Muralidhara *et al*., 2021; Belda-Palazon *et al*., 2022; Henninger *et al*., 2022; Avidan *et al*., 2023). In the second method, Safi and colleagues also express an optimized SnRK1.1 phosphorylation recognition motif (denominated AMPK substrate peptide (ASP)) tagged with GFP, but also fused to a homo-oligomerization coiled-coil sequence (HOTag) (Safi *et al*., 2023). In this design (denominated ASP-SPARK), the phosphorylation of the ASP sequence promotes the oligomerization of the HOTag peptide, inducing a phase separation of this oligomeric reporter peptide. This process produces the appearance of large phase-separated ASP-SPARK condensates visualized as bright fluorescent puncta. Therefore, the quantification of the number of fluorescent puncta by confocal microscopy is the readout of SnRK1.1. kinase activity (Safi *et al*., 2023; Persyn *et al*., 2024). Although the readout ASP-SPARK provides information on SnRK1 at the cellular level, the phase separation principle underlying the technique partially obscures its precise/actual intracellular localization. Additionally, the liquid-liquid phase separation kinetics may limit the ability to accurately determine the temporal dynamics of a SnRK1.1-mediated response.

To complement current methods and overcome some of their limitations in evaluating SnRK1.1 activity, we designed a protocol to quantify the intracellular localization of SnRK1.1. Our protocol, based on the analysis of z-stack sets of images of cells expressing SnRK1.1– eGFP under a native promoter, provides an excellent tool to track changes in the intracellular distribution of SnRK1.1 fractions. The readout of the method is an index between the nuclear and non-nuclear SnRK1.1 fractions, which can be used as a proxy for cell energy status changes and the evaluation of the energy homeostasis response to external or developmental cues. As proof of concept, we evaluated the SnRK1.1-mediated response to DCMU treatments via intracellular distribution as well as T-loop phosphorylation and kinase content. To sum up, this method and the output index provide a new tool to evaluate in a spatiotemporal manner the SnRK1.1 behaviour in response to changes in cell energy levels.

## Materials and methods

### Plant growth conditions

*Arabidopsis thaliana* plants were grown in a growth chamber under a 16 h light/8 h dark cycle, with an irradiance of 120 μmol photon m^−2^ s^−1^ and temperatures of 23 °C/21 °C.

Arabidopsis lines were initially selected on half-strength Murashige and Skoog (1/2 MS) medium supplemented with 35 μg ml^−1^ kanamycin for SnRK1.1–eGFP transgenic lines. Fifteen-day-old plants were used for experiments.

### Cell energy perturbation treatment

Cell energy perturbation experiments were conducted in transgenic Arabidopsis lines expressing SnRK1.1-eGFP (SnRK1.1-*OE*). Two-week-old plants were sprayed with mock solution (dilution of DMSO) and 50 μM DCMU [3-(3,4-dichlorophenyl)-1,1-dimethylurea], each for its designated time before imaging.

### Fluorescence microscopy imaging

Leaves of *A. thaliana* SnRK1.1-*OE* lines were analysed by Laser Scanning Confocal Microscopy (LSCM) using a Zeiss LSM 880 microscope. A Plan-Apochromat 20x/0.8 M27 objective was used for imaging. eGFP was excited with a 488 nm laser and the emission was collected at 490-526 nm. Autofluorescence of chlorophyll was detected in the eGFP channel, using excitation/ emission wavelengths of 543 nm and 690-710 nm, respectively.

### Image analysis

Z-stack images were analyzed using the open source software Fiji/ImageJ (Schindelin *et al*., 2012) and IMARIS (Bitplane) software. In Fiji, images were processed using threshold and Gaussian Blur filter tools to generate nuclei and ER masks. The average intensity fluorescence measurements were obtained from the Measure tool of the ROI manager. Images were pre-processed using ROF filters (Fiji) for IMARIS analyses. Contrast was calculated as (Ir-Ib)/Ib, where Ir and Ib represent the mean intensity of the region of interest and the background, respectively. Then, Nuclei and ER were segmented using the IMARIS automatic surface rendering mode, selecting only those planes that included each region. These software were also used to calculate the mean intensity of both structures for each image.

### Western blot analysis

*Arabidopsis thaliana* transgenic and wild-type lines were used to obtain soluble proteins. Briefly, 200 mg of leaves were ground in liquid nitrogen and homogenized in 4 ml of extraction buffer (50 mM Tris–HCl, pH 7.5; 0.33 M sucrose; 5 mM EDTA; 150 mM NaCl, and 1X complete protease inhibitor cocktail from Roche). The total protein extract was obtained by centrifugation at 10,000 g for 1 min. Proteins were solved by 12% (v/v) SDS– polyacrylamide gel electrophoresis and transferred to a nitrocellulose membrane. Membranes were blocked with 5% (w/v) milk in TBS and then incubated with specific antibodies: anti-GFP-specific (1:3000, Cell Signaling), anti-SnRK1.1 (1:1000, Agrisera), and anti-Tloop (1:1000, Cell Signaling). The membranes were further incubated with anti-rabbit immunoglobulin G (IgG) conjugated with horseradish peroxidase (HRP). Chemiluminescence detection was performed using the SuperSignal West Femto Maximum Sensitivity Substrate (Thermo Scientific), and the signal was visualized using an Amersham Imager 600.

### Photosynthetic measurements

The effects of the DCMU treatments in Arabidopsis plants were measured in 2-week-old control (Col-0) and SnRK1.1-*OE* lines growing at 120 μmol photon m^−2^ s^−1^. Plants from both genotypes were randomly assigned to receive either the treatment (DCMU) or mock (dilution of DCMU solvent, DMSO, in water), by the same protocol as in energy perturbation treatments above. After a specific period of time - 0, 0.25, 1, 2 or 5 hours - PAM photosynthetic measurements and OJIP curves were obtained using FluorPen FP 110 and PhotosynQs MultispeQ PAM fluorometers. For maximum quantum yield of PSII (*F_v_/F_m_*) measurements, leaves were dark-adapted previously for at least 20 minutes, to allow full reoxidation of the electron transport chain.

### Statistical analysis

The N/ER index and flowering time differences between lines were detected using analysis of variance (ANOVA) followed by Tukey’s HSD test (significance threshold p < 0.05), in InfoStat software version 2020e and its interface with R (Di Rienzo *et al*., 2011). N/ER index was obtained using the average fluorescence of each fraction (denominated FL-Nuc-ROI and FL-ER-ROI for nuclear and ER-associated SnRK1.1 fractions, respectively) from z-stacks at each time point (n= 3-4). Flowering time was obtained by averaging the number of days from germination until flower bud observation (0.5 cm) for each line (n = 17). Number of rosette leaves was counted right after bud formation for each line (n= 17). Linear regression of the NER indexes data obtained with the different software (Fiji/IMARIS) was calculated using InfoStat software and setting the y-intercept value to zero. Model significance level p-value <0.001.

## Results

### SnRK1.1 is distributed in the whole volume of pavement cells in different intracellular fractions

Recently, our group and others have revealed that Arabidopsis SnRK1.1 is dually distributed between the surface of the endoplasmic reticulum (ER) and the nucleus (Jamsheer *et al*., 2018; Blanco *et al*., 2019; Gutierrez-Beltran and Crespo 2022). In those works, the distribution of SnRK1.1 was determined in different tissues and plant species by the imaging of a fusion protein with a fluorescent tag (e.g., GFP and YFP). For instance, we used a fusion of a genomic fragment containing SnRK1.1 and its promoter with eGFP at its C-terminal including a linker of 15 amino acids between the last exon and the fluorescent tag to produce transgenic stable lines (SnRK1.1-*OE*s) (Blanco *et al*., 2019). Moreover, we found that SnRK1.1 intracellular distribution was affected by changes in cell energy status produced by inhibition of photosynthesis (Blanco *et al*., 2019).

To conclusively evaluate any change in SnRK1.1 intracellular localization, we developed a new protocol for quantification of its distribution in stable transgenic SnRK1.1-*OE* Arabidopsis lines. First, we confirmed the expression of functional SnRK1.1-eGFP by different approaches (Fig. 1). None of the SnRK1.1-*OE* lines (SnRK1.1-*OE1* and SnRK1.1-*OE2*) exhibited phenotypic differences under long-day conditions, either in soil or in 0.5× MS medium, compared to the control lines (Col-0), transgenic lines expressing eGFP targeted to the nucleus or *snrk1.1*^-/-^ (Fig. 1A). In the initial screening, SnRK1.1-eGFP fusion proteins were detected in leaf tissue of transgenic lines by western blot using commercial antibodies against eGFP (Fig. 1B). Considering that a fraction of SnRK1.1 is attached to the ER, which possesses different subdomains in pavement epidermal cells, we conducted imaging of leaf sections of SnRK1.1-*OE* plants along the z-axis, namely z-stacks. Indeed, the obtained focal planes in each z-stack showed different SnRK1.1 populations ranging from those linked to the cortical ER at the abaxial periclinal cell surface to other fractions localized at the anticlinal ER domain. A set of images spanning from abaxial to adaxial leaf section side belonging to transgenic line SnRK1.1-*OE2* is shown in Fig. 1C. Imaging experiments can be performed either using a single fluorescent channel corresponding to the fluorescent tag of the fusion protein or, alternatively, using multiple channels—including brightfield, an ER marker, or chlorophyll autofluorescence—as references to identify the different regions of the leaf pavement cells (Supplementary Fig. S1). In Fig. 1C, a 3D-rendered visualization of the SnRK1.1 intracellular distribution is shown in a 5-cell ROI of a leaf section, of fluorescence tagged lines expressing SnRK1.1-eGFP. Beyond detecting SnRK1.1-eGFP by western blot (see also Fig. 3) and LSCM images, we assessed the functionality of the fusion protein via flowering time studies. SnRK1.1 overexpression has shown a delayed flowering (Baena-Gonzalez *et al*., 2007; Williams *et al*., 2014; Jeong *et al*., 2015; Belda-Palazon *et al*., 2020). Accordingly, the SnRK1.1-*OE* lines had a 3-day delay in flowering bud formation compared to Col-0 and *snrk1.1^-/-^* lines, confirming the overexpression of an active SnRK1.1-eGFP fusion (Supplementary Table S1, Supplementary Fig. S2A).

**Figure 1.**
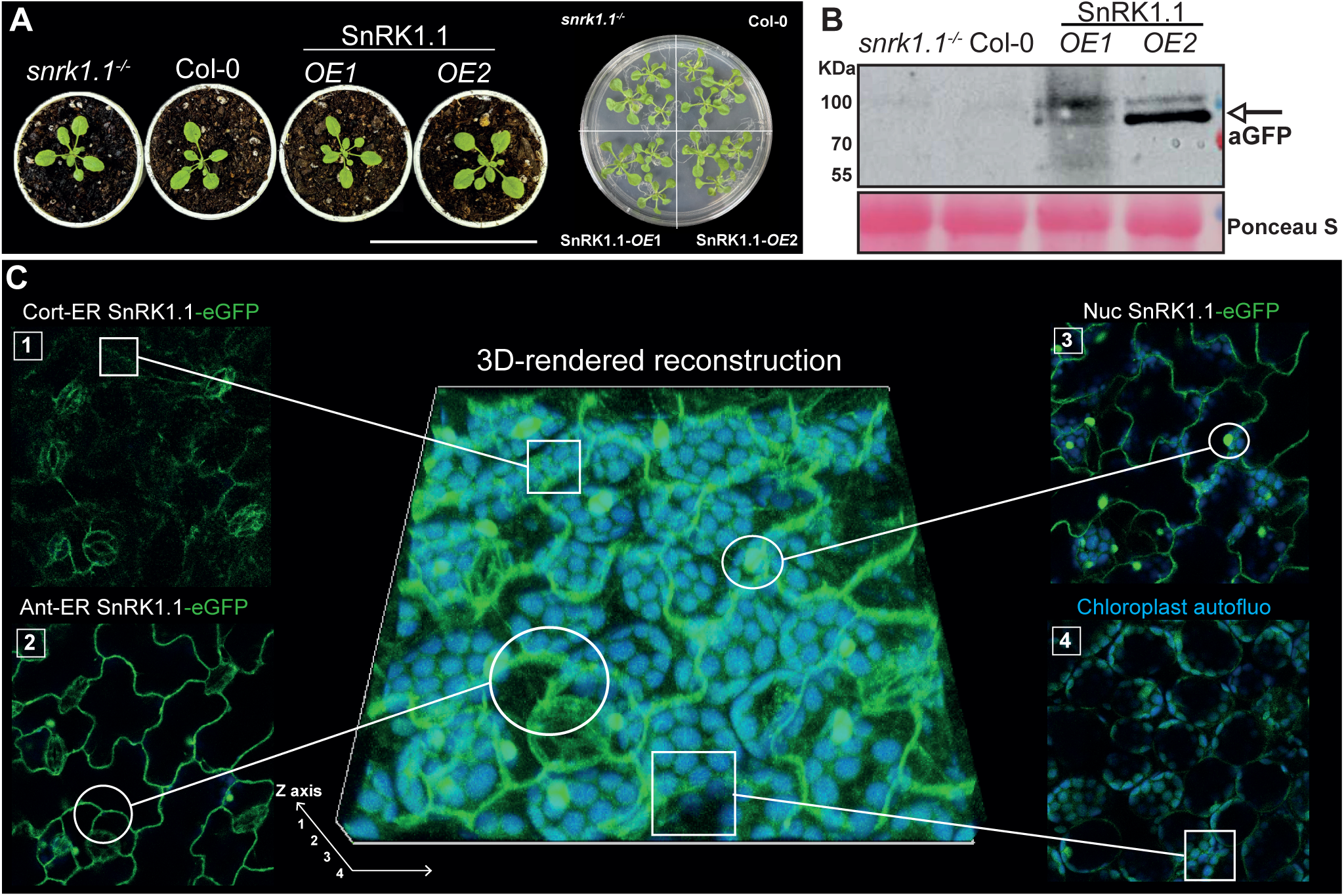
SnRK1.1 is distributed throughout the entire plant cell volume. Set of lines used in this study and their control lines (snrk1.1-/-is the KO line snrk1α1-3 (GABI_579E09), Belda-Palazon et al., 2020) growing under long-day conditions, either in soil or in 0.5× MS medium (A). Western blot of total protein extract of leaf sections from different SnRK1.1-OE plants using anti-GFP antibodies during the initial screening of positive transformants, with image of loading control using Ponceau S (B).Images obtained by LSCM of leaf sections from A. thaliana plants overexpressing SnRK1.1-eGFP (C). A series of images from a z-stack, spanning from the adaxial to the abaxial side: (1) SnRK1.1-eGFP signal in the cortical ER (Cort-ER), (2) SnRK1.1-eGFP signal in the anticlinal ER (Ant-ER), (3) SnRK1.1-eGFP signal in the nucleus (Nuc), and (4) SnRK1.1-eGFP signal in the abaxial-side ER and chloroplasts in the bottom spongy tissue layer. A 3D reconstruction of SnRK1.1 distribution in a leaf pavement cell based on the z-stack images in a plant expressing SnRK1.1-eGFP (Blanco et al., 2019). Chloroplasts are shown in blue, SnRK1.1 in green, and the z-axis/total volume in a white-lined box. Images were captured and processed using Zeiss ZEN 3.10 Lite software. Scale bars: 10 cm (A), 10 µm (C).

After establishing the imaging methodology and plant model, we analyzed multiple z-stacks to develop a quantification protocol of SnRK1.1 intracellular distribution. We chose the comparison of the median fluorescence intensity of the SnRK1.1 fractions at the anticlinal ER domain and in the nucleus as quantification strategy (Fig. 2B). This approach assures consistency in the fluorescence intensity along the SnRK1.1 ER fractions of different z-stacks. The selection of the SnRK1.1 fractions associated with the anticlinal ER domain is representative of the average intensity of the whole non-nuclear fraction of SnRK1.1. Furthermore, the selected fractions have a more even distribution of the SnRK1.1-eGFP signal, avoiding the inclusion of fluorescent foci that are more common in cortical sections. This prevents quantification processes from being skewed by small areas with saturated fluorescence intensities. Additionally, at the cellular level, the SnRK1.1 fraction in the anticlinal ER domain adjacent to the nucleus is more likely to participate directly in nuclear-directed SnRK1-mediated responses than fractions in other ER regions. This provides a more relevant basis for assessing the regulation of nuclear gene expression by SnRK1.

**Figure 2.**
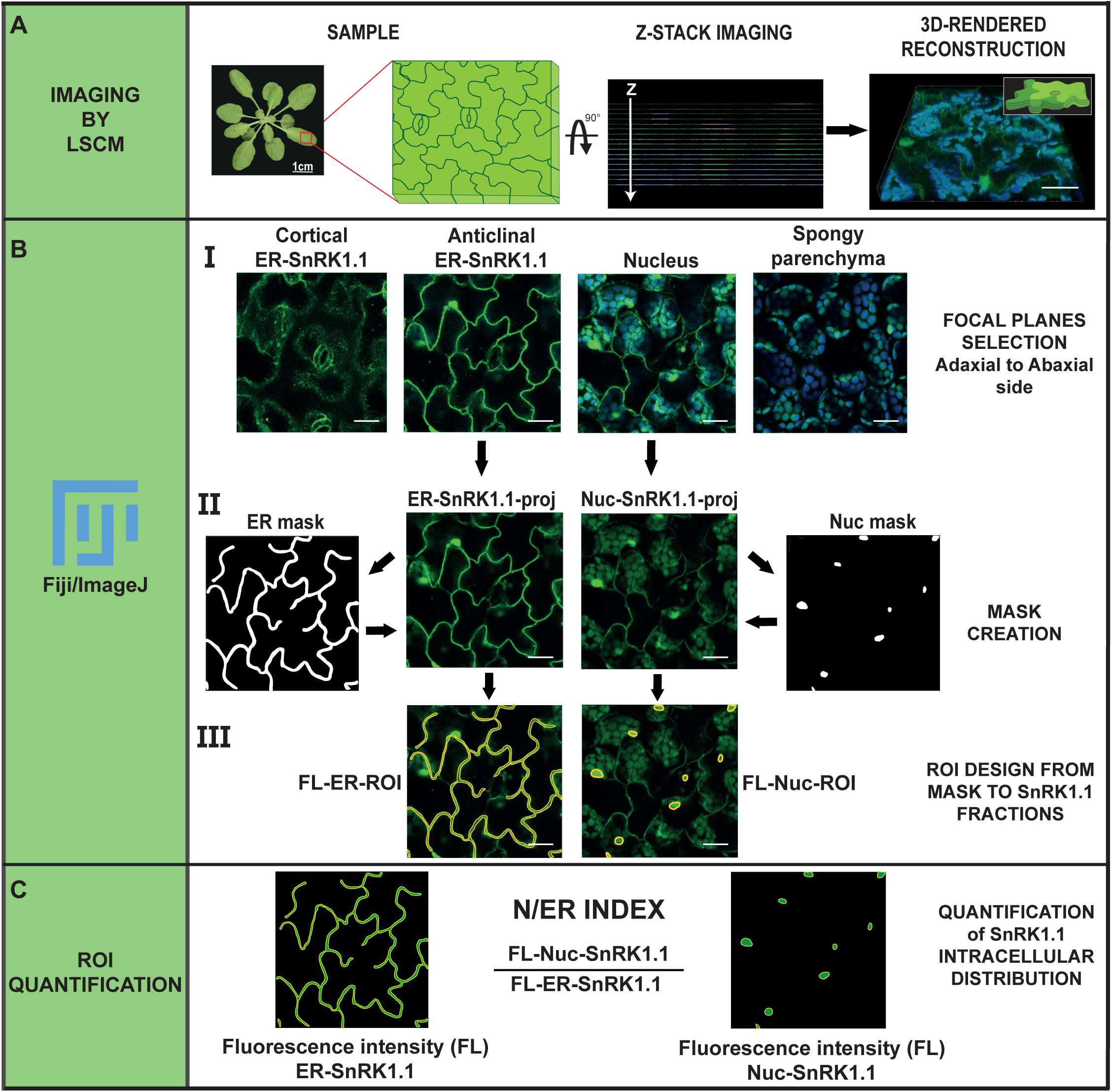
Pipeline of the protocol designed in Fiji/ImageJ to quantify SnRK1.1 intracellular distribution. The protocol is divided into three main stages: Imaging (A), quantification of segmented fluorescence in different SnRK1.1 fractions (B), and parameterization of SnRK1.1 distribution using the Nuc/ER Index (C). A detailed description of each step is provided in the text.

### Quantification of SnRK1.1 fractions and generation of a Nuclear/ER index as concept of cell energy status determinations

Visualizing SnRK1.1 intracellular distribution in z-stacks of leaf sections from SnRK1.1-*OE* plants and selecting focal planes corresponding to SnRK1.1 fractions at the anticlinal ER domain and in the nucleus constitute the first stage of our quantification protocol (Fig. 2A). Using this criterion and measuring strategy, we identified and selected the positions of the focal planes for each SnRK1.1 fraction (ER-associated, hereinafter referred to as ER-SnRK1.1; and nuclear, Nuc-SnRK1.1) in the z-stack of images acquired from sections of a plant leaf (Fig. 2B-I). Any difficulty in identifying the respective focal planes can be overcome using the corresponding image from the brightfield channel and/or from the ER-marker channels of the acquired stacks as a reference. The following steps in the pipeline involved the use of Fiji/ImageJ, an open-source imaging analysis software, and yielded the mean fluorescence intensity values of representative images for each of the SnRK1.1 fractions (Fig. 2B) (Schindelin *et al*., 2012). The z-stack files were opened with Fiji and split into their different channels (use the command *Image > Color > Split Channels*), and sub-stacks corresponding to the SnRK1.1 channel were processed to define masks for segmenting the fluorescence signal of the ER-SnRK1.1 and Nuc-SnRK1.1 (Fig. 2B-I). Three to four z-slices spanning ∼ 4 µm in the z-axis, depending on the acquisition resolution, were selected from each stack, corresponding to ER-SnRK1.1 and Nuc-SnRK1.1 (using the command *Duplicate > Range* and choose z-slice range for different SnRK1.1 fraction) (Fig. 2B-II). Each set of z-slices was then processed to generate a single representative image for the SnRK1.1 fraction by applying the Z-projection function with the average intensity tool (set of commands *Image > Stack > Z-projection > Average Intensity*). Each resulting “projection” image was duplicated for further analysis. These newly generated z-projection images were named “ER-SnRK1.1-proj” and “Nuc-SnRK1.1-proj” (Fig. 2B-II), and served to determine the mean fluorescence intensity for each SnRK1.1 fraction. From these images, two binary images were generated to segment each SnRK1.1 fraction (sequences of commands *Image > Type > 8-bit > Adjust > Threshold > Apply > Process > Filters > Gaussian Blur > Binary > Make Binary > Convert to Mask*). The mask creation process can be automated using the “Auto” tool within the threshold settings for segmentation correction and may be executed in batch mode by recording the steps and creating a macro (Plugins > Macros > Record). Alternatively, ER-marker brightfield images can be used to refine the generated masks, which can improve the accuracy of SnRK1.1 intensity quantification in the “ER-SnRK1.1-proj” and “Nuc-SnRK1.1-proj” images.

Next, a region of interest (ROI) was defined for each projection image using the corresponding masks, which yielded the average fluorescence of the ER- and Nuc-SnRK1.1 fractions. The ROI was generated using the Fiji Wand tool within ROI manager by clicking on each mask. If the mask creation process yielded multiple ROIs, they were combined using the command ROI management > More > OR (combine) > Add. The mean fluorescence intensity within the ROIs of ER-SnRK1.1 and Nuc-SnRK1.1 was then measured (ROI management > Select ROIs > Measure > Mean Intensity), resulting in two values: “FL-ER-ROI” and “FL-Nuc-ROI” (Fig. 2B-III). Finally, the ratio between “FL-Nuc-ROI” and “FL-ER-ROI” was calculated, yielding a numerical value referred to as “N/ER index” (Fig. 2C). This N/ER index quantifies the relative distribution of SnRK1.1 between the ER-associated and nuclear fractions within the cell volume, providing insight into its intracellular localization and dynamic behaviour.

### Proof-of-Concept: Evaluating the Effect of DCMU on SnRK1.1 Intracellular Distribution

Changes in the distribution of SnRK1.1 have been observed during developmental processes and in response to various environmental cues (Zhai *et al*., 2017; Blanco *et al*., 2019; Han *et al*., 2020; Belda-Palazon *et al*., 2022; Shi *et al*., 2024). To validate our protocol, we evaluated the previously reported effect of photosynthetic electron transport blockage on the intracellular distribution of SnRK1 (Blanco *et al*., 2019). Our previous results showed that changes in SnRK1.1 localization primarily occurred in vascular tissues after 1 hour of treatment with DCMU, an inhibitor that binds to the quinone binding site B (Q_B_) of photosystem II and blocks electron flow from PSII to plastoquinone (Takahashi *et al*., 2010) (Supplementary Fig. S3A and Table S2).

Using 2-week-old SnRK1.1-*OE* plants, we imaged leaf sections from plants sprayed with 50 µM DCMU or mock-treated over time (Fig. 3). In pavement cells expressing SnRK1.1-eGFP, we observed the same behaviour of the fusion protein previously reported in vascular tissue (Fig. 3). An apparent increase in the Nuc-SnRK1.1 fraction was observed at the expense of the ER-SnRK1.1 fraction, as visualized in 3D reconstructions of the analyzed sections (Fig. 3A). For each time point, we applied our protocol to at least three z-stack series of various individuals, across three independent experiments (Fig. 3B, Supplementary Fig. S4). The visual trend observed in LSCM images was confirmed through quantification, which showed a statistically significant increase in the N/ER index between 1 and 5 hours after treatment (*i.e*., from 2,38 ± 0.38 before treatment to 2.76 ± 0.65 and 3.21 ± 0.21, after 1 and 5 hs after treatment, respectively, p<0.01) (Fig. 3A-B, see also Bottom table Fig. 4).

**Figure 3.**
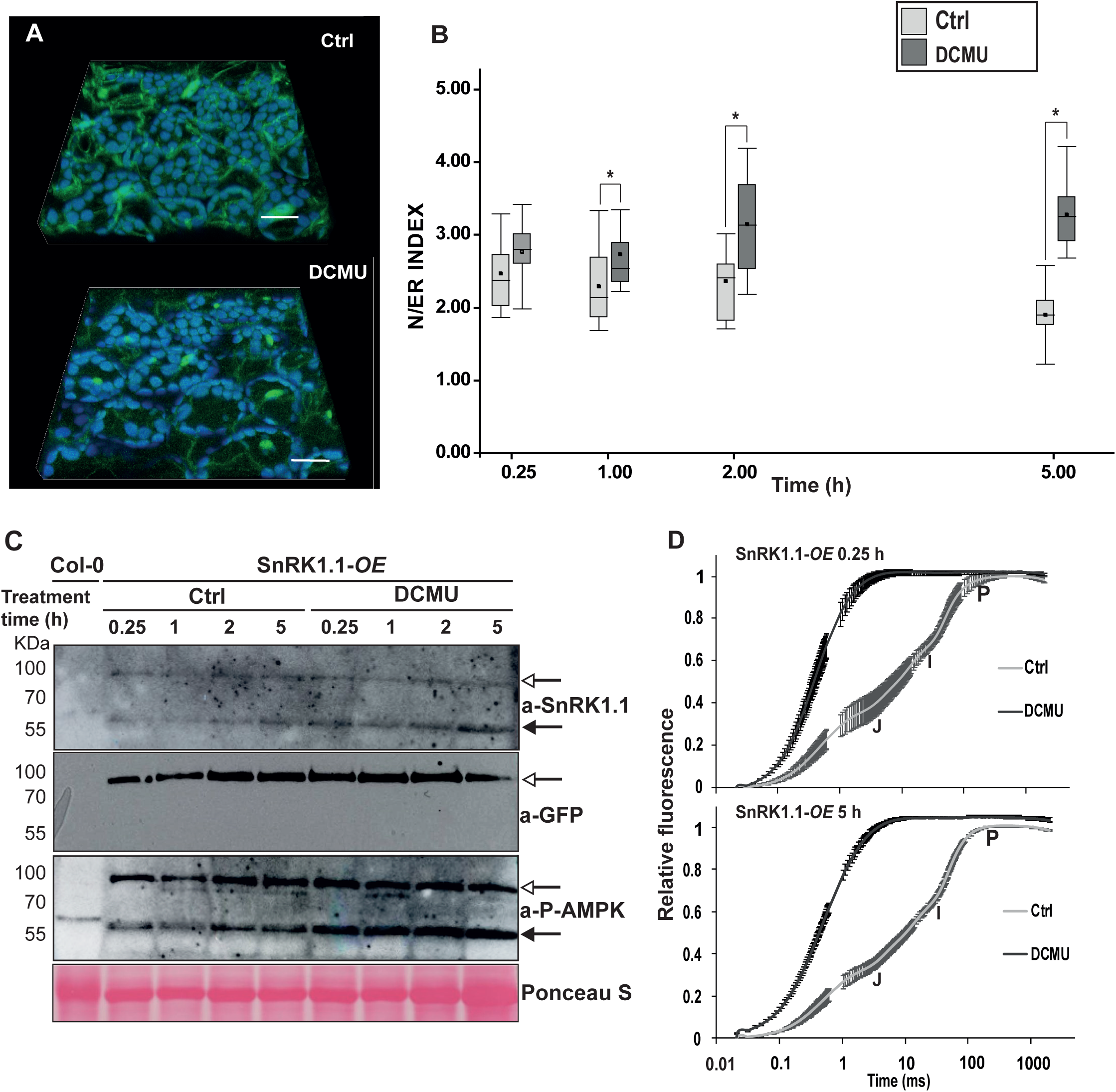
Study of the SnRK1.1-mediated response to the blockage of photosynthesis. An integrative analysis of SnRK1.1 intracellular dynamics was performed using the N/ER index quantification protocol to understand its response to photosynthesis blockage. A 3D reconstruction of SnRK1.1-eGFP distribution changes in SnRK1.1-*OE* plants treated with DCMU for 2 hs (A). The N/ER index was calculated from z-stacks of DCMU- (dark grey box plots) and mock-treated (Ctrl, light grey box plots) plants (B). In parallel, endogenous and alien fusion protein levels were assessed by Western blot using anti-SnRK1.1 (Agrisera), phospho-AMPKα (T172) (Cell Signaling), and GFP (Agrisera) antibodies (C). The signal corresponding to the exogenous fusion protein is indicated with an open-headed arrow, while the endogenous SnRK1.1 signal is marked with a solid-headed arrow. The effectiveness and stability of DCMU treatment over time were confirmed by chlorophyll a fluorescence transient in OJIP curves (D). The DCMU treatment abolished the JIP phases of the OJIP curves (solid symbols, DCMU curves) compared to mock-treated plants (open symbols, Ctrl curves) throughout the evaluated period, as represented by measurements taken at 15 minutes (upper graph) and 5 hours (lower graph) indicating a total stop in photosynthetic energy production.

**Figure 4.**
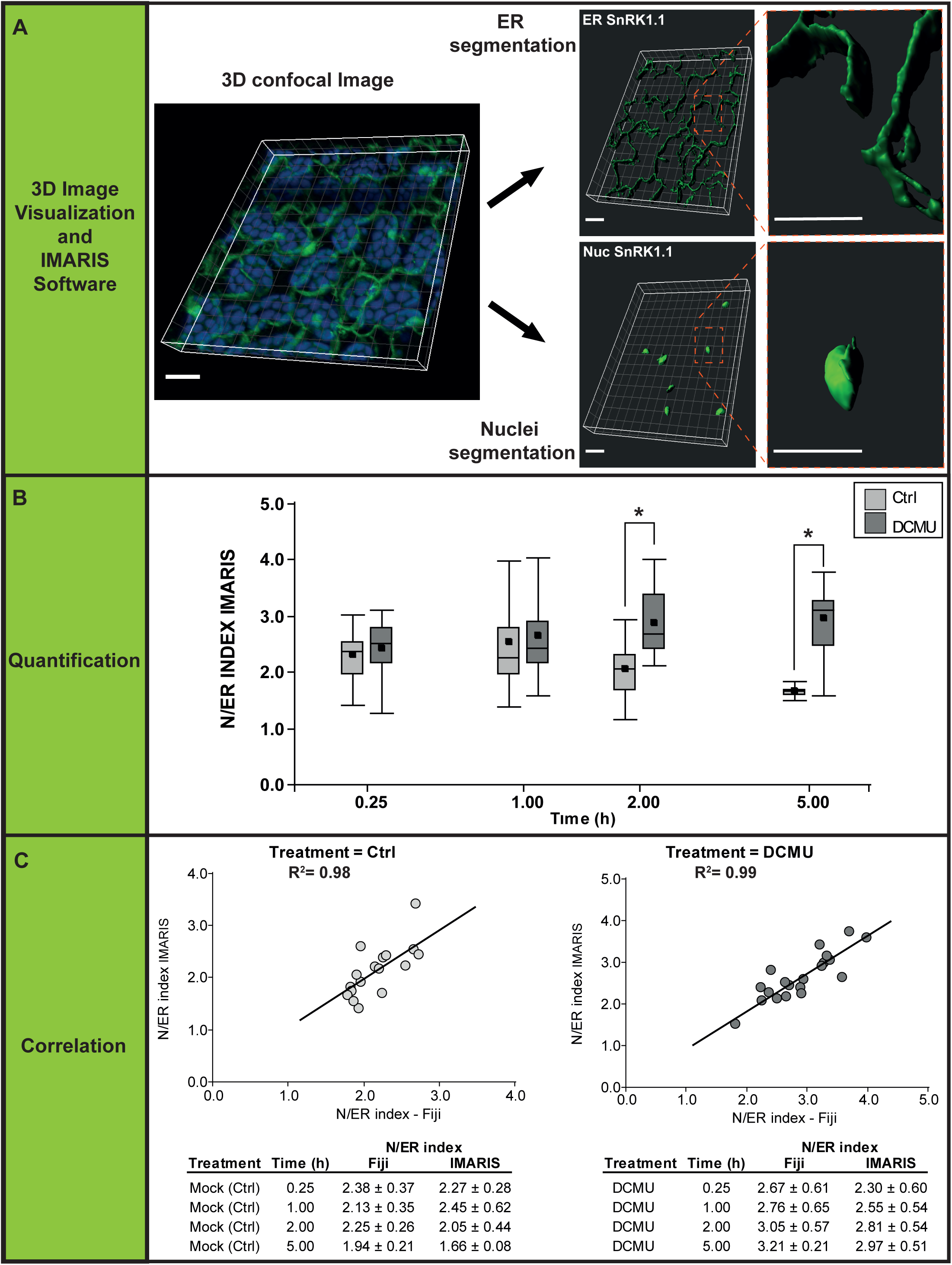
Quantification of N/ER index protocol using an alternative segmentation software. Representative 3D image reconstruction and segmentation obtained from SnRK1.1-eGFP signal in Nuclei and ER using a commercial 3D specialized microscopy analysis software (IMARIS). Scale bar = 20 μm. **(A)**. Analysis of SnRK1.1 distribution over time (0 to 5h) using the N/ER index calculated with IMARIS software, for mock-treated (Ctrl, light-grey) or 50 μM DCMU-treated plants (dark-grey). Differences between treatments were detected using one-way ANOVA followed by Tukey HSD test (p-value < 0.05) **(B)**. Scatter-plot for pairs of N/ER indexes obtained with Fiji or IMARIS (same set of confocal images used in the calculation of the Nuclear and ER sections of SnRK1.1), for mock-treated (Ctrl, light-grey) or DCMU-treated plants (dark-grey). Linear regression was used to estimate the correlation coefficient for each experiment (R2). Table of average N/ER index obtained with Fiji and IMARIS (mean ± S.D., n=4-7) **(C)**.

To gain further insight into this response, we evaluated endogenous and overexpressed SnRK1.1 protein levels by western blot on imaged sections (Fig. 3C). No significant changes were observed in either endogenous SnRK1.1 or SnRK1.1-eGFP content in DCMU- or mock-treated leaf sections (Fig. 3C upper blot). This was further confirmed using anti-GFP antibodies, which specifically recognize SnRK1.1-eGFP (Fig. 3C medium blot). The absence of high-mobility signals in the 20–35 kDa range confirmed the integrity of the fusion protein, indicating that the fluorescence signal identified as Nuc-SnRK1.1 (Fig. 2B) accurately corresponds to SnRK1.1-eGFP, rather than to a potentially cleaved eGFP tag relocalized to the nucleus. Furthermore, we evaluated whether phosphorylation of the T-loop was associated with these changes in SnRK1.1 intracellular distribution. As observed for SnRK1.1 protein levels, no significant differences in T-loop phosphorylation were detected in either endogenous SnRK1.1 or SnRK1.1-eGFP in response to DCMU treatment (Fig. 3C lower blot). Finally, using various photosynthetic measurements, including OJIP curves, we confirmed that DCMU treatment resulted in a complete and sustained blockage of photosynthetic electron transport throughout the experiment (Fig. 3D, Supplementary Fig. S3 and S5 and Table S2). Analysis of the first and last time points of the treatment showed the abolishment of the JIP phases, indicating an effective blockage of the electron flow downstream of Q_A_ in Photosystem II (PSII) (upper and lower panels in Fig. 3D; full timeframe in Supp. Fig. S5) (Tóth et al., 2005).

The drop in *q_L_* values to ∼0,2, and the complete loss of the Y(II) parameter confirm the absence of photosynthetic transport in DCMU-sprayed leaves throughout the treatment period, regardless of SnRK1.1-eGFP overexpression (Supplementary Fig. S3). The DCMU treatment caused a small decrease in the maximum quantum yield (*F_v_*/*F_m_*), which might indicate a perturbation in the funnelling of absorbed energy into PSII, but not permanent damage.

These results demonstrate the changes in SnRK1.1. distribution, quantified by the N/ER index, were triggered by a total interruption of photosynthetic electron flow and the resulting decrease in photosynthetic energy production over the 5-hour treatment period.

### Compatibility of our quantification protocol for SnRK1.1 intracellular distribution with other software tools

To assess the compatibility of our quantification protocol, which includes various segmentation and quantification steps, with alternative analysis tools, we analyzed the DCMU treatment data using IMARIS (Bitplane, Belfast, Northern Ireland, UK). This software is widely used for advanced multidimensional image visualization, segmentation, and quantitative analysis (Fig. 4). Specifically, we used the same z-stack sets as in Fig. 3B from mock- and DCMU-treated plants over time (0.25-5h), to calculate the N/ER index based on the fluorescent intensity in different subcellular fractions of SnRK1.1-eGFP, by segmentation with IMARIS. Before segmentation, the images were pre-processed with a ROF filter (Fiji), which reduces noise while preserving edges and enhancing image quality. Next, nuclei and ER were segmented using IMARIŚs automatic surface rendering mode, applying a user-defined intensity threshold to identify structures with a contrast value above ∼3, and a size threshold of 8 µm for nuclei segmentation, set to exclude small, non-biologically relevant elements. Only those planes corresponding to each region of interest were selected for analysis (Fig. 4A). Finally, the software was used to calculate the mean intensity of both structures in each segmented 3D image. As in Fiji-based analysis, we determined SnRK1.1 changes in its intracellular distribution along time by calculating the N/ER index for mock- (Ctrl) and DCMU-treated plants (Fig. 4B). Consistent with our Fiji procedure (Fig. 3B), we observed an enrichment of SnRK1.1 in the nucleus after 2 and 5 hours of DCMU-treated plants (Fig. 4B).

The scatter-plot for the pair of N/ER indexes obtained by our protocol and by using IMARIS showed a linear correlation (Fig. 4C, Supplementary Table S3), with highly significant correlation coefficients of r^2^=0.98, (p < 0.0001, mock) and r^2^=0.99 (p < 0.0001, DCMU), respectively. The high correlation was also observed in the averaged values per time-point (Fig. 4C, Supplementary Table S3). To estimate the robustness and precision for both protocols, we calculated the coefficient of variation throughout the complete set of N/ER indexes (n=38 and n=45, for mock and DCMU, respectively). Coefficients of variation (CVs) for Fiji measurements were between 12.6% ± 6.1% and 17.0% ± 8.4% for mock- and DCMU-treated plants, respectively. For IMARIS, CVs were between 15.9% ± 9.6% and 19.3% ± 8.1% for mock- and DCMU treated-plants, respectively. CVs maintained moderate across different experiments (mock and DCMU), with a slight overperformance for Fiji measurements. Overall, this evaluation confirmed that the quantification protocol is robust, reliable and compatible with different segmentation and quantification approaches.

## Discussion

Significant efforts have been made to understand how SnRK1 restores energy homeostasis since the initial characterization of Arabidopsis *SnRK1.1* mutant lines (Baena-Gonzalez *et al*., 2007). Beyond identifying its target genes, phosphorylation consensus motifs, target proteins, and interactors, the recent discovery of distinct intracellular fractions of SnRK1.1 has provided new insights into its activity (Jamsheer *et al*., 2018a; Blanco *et al*., 2019). Furthermore, changes in SnRK1.1 intracellular localization in response to the blockage of photosynthesis by DCMU have also contributed to deciphering its activation mechanisms (Fig. 3) (Blanco *et al*., 2019; Ramon *et al*., 2019). Currently, there is consensus that the activity of SnRK1.1 signalling pathway is linked with its spatiotemporal fingerprint, a characteristic also observed in its opisthokont counterparts (Zong *et al*., 2019; Gutierrez-Beltran and Crespo 2022). This emerging model is built upon the idea of a spatial separation of roles for SnRK1.1: sensing energy imbalance at the ER and mediating metabolic reprogramming by regulating gene expression in the nucleus to restore energy homeostasis (Wurzinger *et al*., 2018; Blanco *et al*., 2019; Crepin and Rolland 2019; Ramon *et al*., 2019; Gutierrez-Beltran and Crespo 2022; Peixoto and Baena-Gonzalez 2022). Due to its central role, this form of "retrograde signalling" also involves intracellular crosstalk with other regulatory pathways, notably its co-regulation with the TOR complex (Nukarinen *et al*., 2016; Belda-Palazon *et al*., 2022; Jamsheer *et al*., 2022).

Based on this model, we designed a protocol to assess SnRK1.1 activity by quantifying its intracellular distribution. Starting from z-stacks of fluorescent images obtained from stable SnRK1.1-eGFP lines, the distribution of GFP signal between the nucleus (Nuc-SnRK1.1) and the ER (ER-SnRK1.1) is quantified using Fiji, which serves as a tool for segmenting and quantifying each fraction separately. The results are expressed through a new parameter, the N/ER index, which normalizes expression differences between cells. The choice of Fiji in this protocol offers a simple, free, and open-source solution for both segmentation and quantification, and it is suitable for automation through Fiji macros. In terms of segmentation, the effectiveness of Fiji is enhanced by two factors: *(i)* the selection of specific plant tissue to image, and *(ii)* the choice of z-planes within the cells. Leaf tissue is an optimal model for evaluating the impact of growth conditions on plant cell energy status and SnRK1.1 distribution (*i*). In particular, the pavement cells selected in our study exhibit photosynthetic performance that is sensitive to external cues (Dopp et al., 2023). Additionally, due to their distinctive morphology, pavement cells facilitate the segmentation of the different compartments where SnRK1.1 is associated, namely nucleus and ER surface *(ii)* (Fig. 2A). For Nuc-SnRK1.1, selecting focal planes at the most abaxial position of the z-stacks, just before the position of spongy parenchyma in the z-axis, enables easier segmentation of nuclei by discrete ROIs, thereby improving the quantification of this fractions. As previously mentioned, the focal planes spanning the SnRK1.1 fraction associated with the anticlinal ER provide a representative intensity value for the entire ER-associated pool. The validity of this segmentation and quantification strategy using Fiji was confirmed by the similar N/ER index results obtained with IMARIS, used here as an alternative segmentation software for quantifying the Nuc-SnRK1.1 and ER-SnRK1.1 intensities (Fig. 4). Moreover, experiments where the plants were previously treated with DCMU indicate no changes in the levels of SnRK1.1 or the SnRK1.1-eGFP fusion protein (Fig. 3C), supporting the conclusion that changes in the N/ER index primarily reflect an increase in the fluorescence signal from the Nuc-SnRK1.1 fraction, at the expense of the ER-SnRK1.1 fraction (Fig. 3 and 4, DCMU treatment). These results are consistent with previous observations in vascular tissue treated with DCMU and ABA-induced response in roots (Blanco *et al*., 2019; Belda-Palazon *et al*., 2022). Interestingly, these changes in SnRK1.1 intracellular distribution also occurred independently of T-loop phosphorylation (Fig. 3C). Structural predictions and *in planta* analyses have also shown that T-loop phosphorylation is stable and may play a limited role in modulating SnRK1.1-mediated response (Emanuelle *et al*., 2016; Ramon *et al*., 2019). To date, significant changes in T-loop phosphorylation have only been observed within 30 minutes following submergence treatment (Cho *et al*., 2016). Although a regulatory role for T-loop phosphorylation cannot be ruled out, our results support the current working model in which SnRK1-mediated responses are largely governed by changes in SnRK1.1 localization (see below, alternative methods for determining SnRK1.1 activity).

One advantage of studying SnRK1.1 intracellular distribution via the N/ER index is the compatibility of this approach with different image segmentation and quantification methods. Both Fiji—the initial method used to establish the protocol—and IMARIS yielded similar N/ER indexes when evaluating SnRK1.1 response to photosynthesis blockage (Figs. 3 and 4), with a single statistical difference observed at 1h of DCMU treatment. A possible explanation is that the automatic thresholding routine used with IMARIS produces the inclusion of a broader population of ER-associated SnRK1.1, including a fraction located near cortical adaxial regions. These areas are characterized by fewer and smaller chloroplasts with lower photosynthetic activity (Mackenzie and Mullineaux 2022). Consequently, a higher ER-SnRK1.1 signal may result from reduced photosynthetic inhibition compared to the anticlinal ER regions adjacent to highly active mesophyll chloroplasts. A similar pattern was previously observed in the dynamic behaviour of SnRK1.1 in vascular tissues (Blanco *et al*., 2019). Despite these minor differences, DCMU treatment effectively blocked photosynthetic electron transport and altered cellular energy status, validating the protocol. The loss of photosynthetic activity was confirmed by OJIP transients and PAM measurements (Fig. 3; Supplementary Figs. S3 and S5). DCMU induced complete inhibition of photosynthetic electron transport in illuminated leaves of both control and SnRK1.1-*OE*, as shown by significant reductions in *q_L_* and Y(II) (Supplementary Fig. S5). The disappearance of characteristic shoulders and plateaus at photosynthesis induction in the OJIP curves further confirmed this effect (Supplementary Fig. S3). A mild and transient reduction in *F_v_*/*F*_m_ was also observed, indicating no lasting damage and ruling out any permanent oxidative stress caused by the DCMU treatment. Importantly, under both DCMU- and mock-treatments, the robustness and reproducibility of our quantification protocol were confirmed by the strong correlation between the results of Fiji and IMARIS.

This was evidenced by the high r² value (∼0.98) and highly significant p values (<0.0001), confirming the reliability of the N/ER index as a comparative metric for SnRK1.1 localization. The homogeneity, stability, and robustness of N/ER index values across segmentation methods, treatments, and timepoints highlight its strength as a reliable parameter to quantify SnRK1.1 intracellular distribution. Under control conditions (long-day 16:8 h, 100–120 µmol m^-2^ s^-1^, ZT3–ZT8), N/ER index value (∼ 2.25) can serve as a proxy for the energy status of non-stressed plant cells. Accordingly, deviations from this index value can be used as readouts for assessing the quality of growth conditions and their effect on cell energy status. Needless to say that more extensive applications of the N/ER index using different growth conditions are needed to confirm it. Independent of this, our approach can be readily integrated with other existing tools to study SnRK1 signalling activity. *A priori*, its compatibility with phase separation-based visualization of kinase activity using ASP-SPARK constructs may not be straightforward (Safi *et al*., 2023). However, ASP-SPARK reporter method could be applied sequentially to determine the kinetics of general SnRK1.1 activity in response to a stimulus, followed by N/ER index quantification to map the spatial localization of downstream phosphorylation events. In parallel, ACC phosphorylation motif-based reporter methods (Sanagi *et al*., 2021), in particular newer versions incorporating an NLS and eGFP for nuclear SnRK1.1 activity quantification (Muralidhara *et al*., 2021; Belda-Palazon *et al*., 2022; Avidan *et al*., 2023), share spatiotemporal compatibility with our N/ER index protocol. When used together, these tools can help dissect the sequence of molecular events regulating SnRK1.1 activation and deactivation during energy homeostasis recovery. These studies can be useful to address three key questions regarding SnRK1.1 activity: *i)* Is SnRK1.1 activity fully independent of T-loop phosphorylation? *ii)*, is T-loop phosphorylation required for SnRK1.1 intracellular localization?, and *iii)* are changes in localization part of the mechanism that deactivates SnRK1.1 or *vice versa*? So far, current evidence supports a positive answer to *(i)*, but the roles of T-loop phosphorylation in localization and regulation of SnRK1.1 activity (*ii* and *iii*) remain unknown. An interesting application of these complementary methods would be the design of an ACC-based reporter driven by SnRK1.1 regulatory sequences and targeted to the ER surface using N-terminal tags such as the N-myristoylation site of SnRK1β1 and β2 or an ER membrane marker sequence (McFarlane *et al*., 2017; Ramon *et al*., 2019; Wang *et al*., 2019).

In summary, the N/ER index offers a simple, rapid, and robust method to quantify SnRK1.1 intracellular dynamics and pathway activity over time. Beyond SnRK1.1, our quantification protocol can be adapted to study any signalling protein whose activity involves changes in intracellular distribution. Accordingly, it serves as a useful tool to refine signalling protein interactomes by assigning spatial and temporal context to protein–protein interactions. Developing specific indexes for different intracellular compartments—and tracking these values over time for a given protein—represents a promising strategy to establish a spatiotemporal hierarchy that enables the differentiation between direct and indirect interactors. Among the most compelling candidates for this type of analysis is the TOR complex, which antagonizes SnRK1 activity and may be particularly well suited for such spatiotemporal studies.

## AUTHOR CONTRIBUTION

NEB conceived the project and experiments. NEB and CB conceptualized N/ER index. DL created plant lines. CB grew the plants, performed microscopy and DCMU experiments, designed the protocol with Fiji and performed the N/ER index acquisition. FJH helped CB with Fiji protocol. CD and VL performed the N/ER index measurements using IMARIS. JB obtained and analyzed the photosynthetic measurements. ADL performed Western Blot. CMF helped obtaining Flowering data, performed statistics and helped with the linear regression analysis. JD helped in discussion. NEB, CB, CMF, CD and VL wrote the manuscript. All authors read and approved the manuscript.

## DATA AVAILABILITY

Data is included in Supplementary information. Any data is also available upon request to.

## Supplementary data

**Supplementary table 1.**
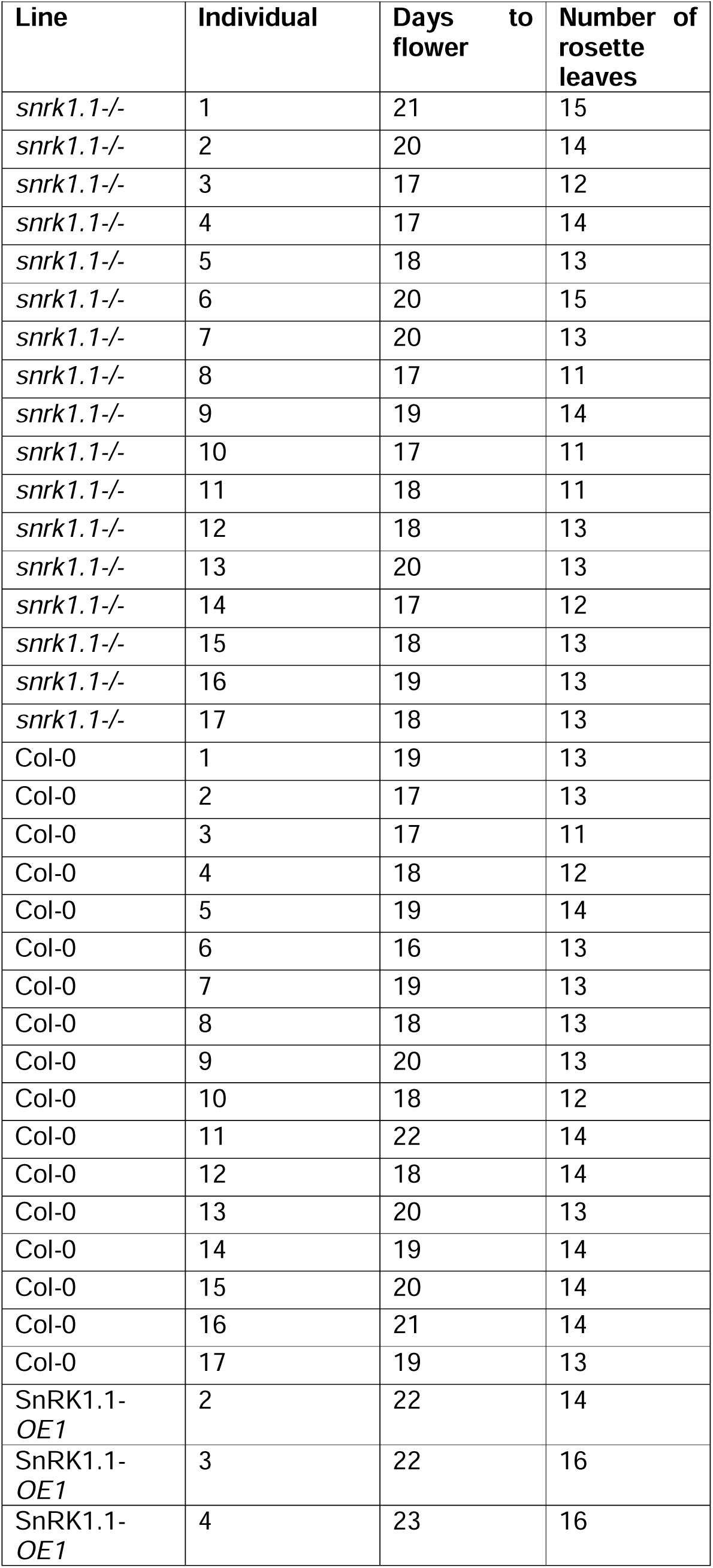
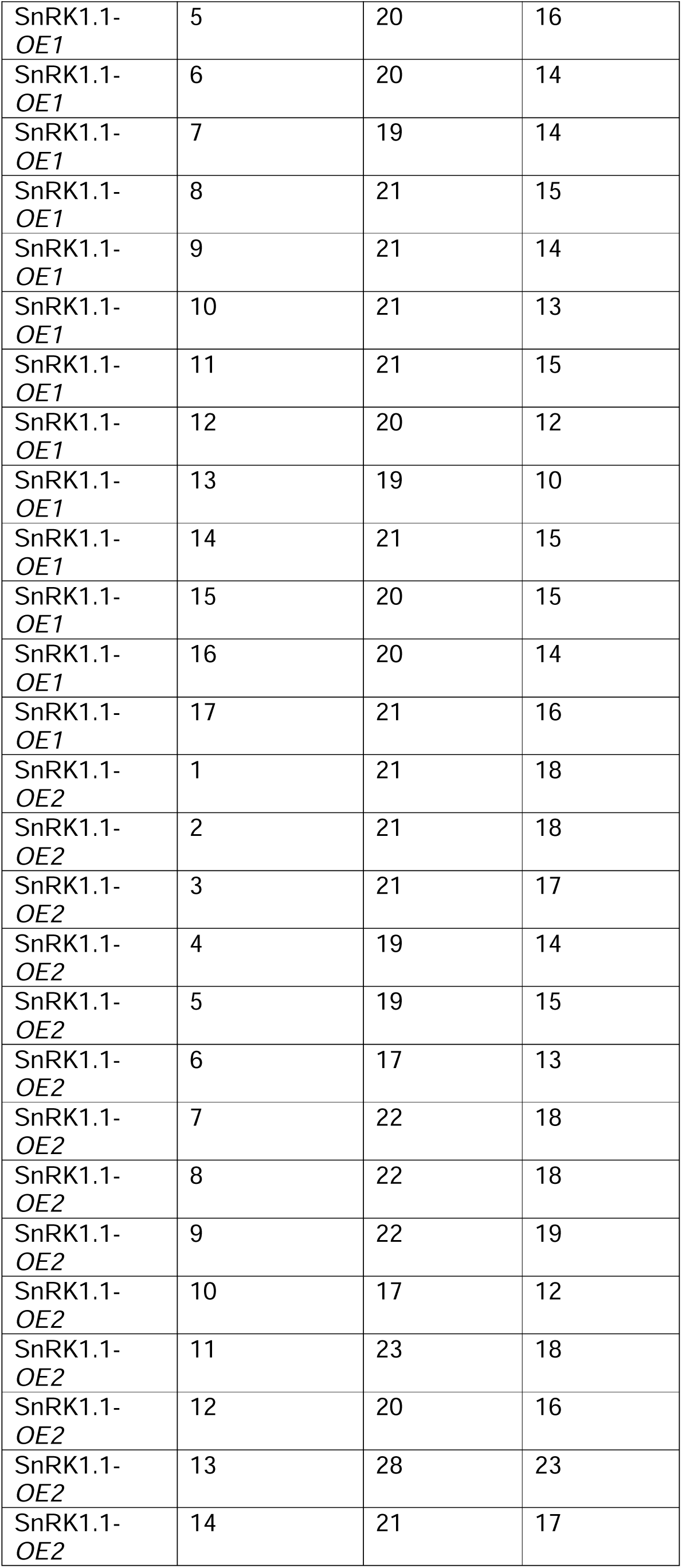
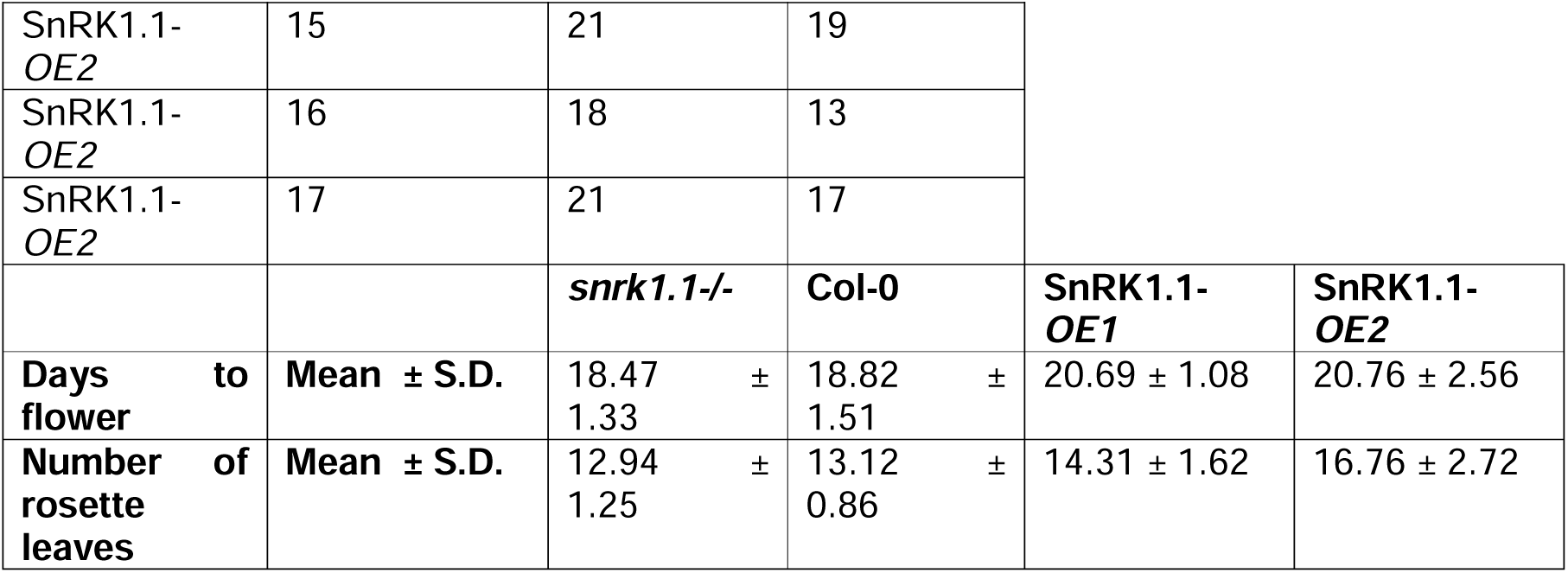
Flowering time in SnRK1.1 lines.

| Supplementary table 1- Flowering time in SnRK1.1 lines |  |  |  |
| --- | --- | --- | --- |
| Line | Individual | Days to flower | Number of rosette leaves |
| *snrk1.1-/-* | 1 | 21 | 15 |
| *snrk1.1-/-* | 2 | 20 | 14 |
| *snrk1.1-/-* | 3 | 17 | 12 |
| *snrk1.1-/-* | 4 | 17 | 14 |
| *snrk1.1-/-* | 5 | 18 | 13 |
| *snrk1.1-/-* | 6 | 20 | 15 |
| *snrk1.1-/-* | 7 | 20 | 13 |
| *snrk1.1-/-* | 8 | 17 | 11 |
| *snrk1.1-/-* | 9 | 19 | 14 |
| *snrk1.1-/-* | 10 | 17 | 11 |
| *snrk1.1-/-* | 11 | 18 | 11 |
| *snrk1.1-/-* | 12 | 18 | 13 |
| *snrk1.1-/-* | 13 | 20 | 13 |
| *snrk1.1-/-* | 14 | 17 | 12 |
| *snrk1.1-/-* | 15 | 18 | 13 |
| *snrk1.1-/-* | 16 | 19 | 13 |
| *snrk1.1-/-* | 17 | 18 | 13 |
| Col-0 | 1 | 19 | 13 |
| Col-0 | 2 | 17 | 13 |
| Col-0 | 3 | 17 | 11 |
| Col-0 | 4 | 18 | 12 |
| Col-0 | 5 | 19 | 14 |
| Col-0 | 6 | 16 | 13 |
| Col-0 | 7 | 19 | 13 |
| Col-0 | 8 | 18 | 13 |
| Col-0 | 9 | 20 | 13 |
| Col-0 | 10 | 18 | 12 |
| Col-0 | 11 | 22 | 14 |
| Col-0 | 12 | 18 | 14 |
| Col-0 | 13 | 20 | 13 |
| Col-0 | 14 | 19 | 14 |
| Col-0 | 15 | 20 | 14 |
| Col-0 | 16 | 21 | 14 |
| Col-0 | 17 | 19 | 13 |
| SnRK1.1-<br>*OE1* | 2 | 22 | 14 |
| SnRK1.1-<br>*OE1* | 3 | 22 | 16 |
| SnRK1.1-<br>*OE1* | 4 | 23 | 16 |
| SnRK1.1-<br>*OE1* | 5 | 20 | 16 |
| SnRK1.1-<br>*OE1* | 6 | 20 | 14 |
| SnRK1.1-<br>*OE1* | 7 | 19 | 14 |
| SnRK1.1-<br>*OE1* | 8 | 21 | 15 |
| SnRK1.1-<br>*OE1* | 9 | 21 | 14 |
| SnRK1.1-<br>*OE1* | 10 | 21 | 13 |
| SnRK1.1-<br>*OE1* | 11 | 21 | 15 |
| SnRK1.1-<br>*OE1* | 12 | 20 | 12 |
| SnRK1.1-<br>*OE1* | 13 | 19 | 10 |
| SnRK1.1-<br>*OE1* | 14 | 21 | 15 |
| SnRK1.1-<br>*OE1* | 15 | 20 | 15 |
| SnRK1.1-<br>*OE1* | 16 | 20 | 14 |
| SnRK1.1-<br>*OE1* | 17 | 21 | 16 |
| SnRK1.1-<br>*OE2* | 1 | 21 | 18 |
| SnRK1.1-<br>*OE2* | 2 | 21 | 18 |
| SnRK1.1-<br>*OE2* | 3 | 21 | 17 |
| SnRK1.1-<br>*OE2* | 4 | 19 | 14 |
| SnRK1.1-<br>*OE2* | 5 | 19 | 15 |
| SnRK1.1-<br>*OE2* | 6 | 17 | 13 |
| SnRK1.1-<br>*OE2* | 7 | 22 | 18 |
| SnRK1.1-<br>*OE2* | 8 | 22 | 18 |
| SnRK1.1-<br>*OE2* | 9 | 22 | 19 |
| SnRK1.1-<br>*OE2* | 10 | 17 | 12 |
| SnRK1.1-<br>*OE2* | 11 | 23 | 18 |
| SnRK1.1-<br>*OE2* | 12 | 20 | 16 |
| SnRK1.1-<br>*OE2* | 13 | 28 | 23 |
| SnRK1.1-<br>*OE2* | 14 | 21 | 17 |

**Supplementary table 1- Flowering time in SnRK1.1 lines**
|  |  |  |  |  |  |
| --- | --- | --- | --- | --- | --- |
| SnRK1.1-<br>*OE2* | 15 | 21 | 19 |  |  |
| SnRK1.1-<br>*OE2* | 16 | 18 | 13 |  |  |
| SnRK1.1-<br>*OE2* | 17 | 21 | 17 |  |  |
|  |  | *snrk1.1*-/- | Col-0 | SnRK1.1-<br>*OE1* | SnRK1.1-<br>*OE2* |
| Days to flower | Mean ± S.D. | 18.47 ± 1.33 | 18.82 ± 1.51 | 20.69 ± 1.08 | 20.76 ± 2.56 |
| Number of rosette leaves | Mean ± S.D. | 12.94 ± 1.25 | 13.12 ± 0.86 | 14.31 ± 1.62 | 16.76 ± 2.72 |

**Supplementary Table S2.** Data of Photosynthetic parameters results.

| Genotype | Treatment | Treatment time (h) | Y(II) | Y(NO) | Y(NPQ) | qL | Fv/Fm |
| --- | --- | --- | --- | --- | --- | --- | --- |
| SnRK1.1-*OE* | Control | 0 | 0.41 ± 0.04 | 0.12 ± 0.01 | 0.47 ± 0.05 | 0.71 ± 0.03 | 0.842 ± 0.005 |
|  |  | 0.25 | 0.44 ± 0.04 | 0.14 ± 0.01 | 0.43 ± 0.05 | 0.66 ± 0.02 | 0.81 ± 0.05 |
|  |  | 1 | 0.42 ± 0.03 | 0.12 ± 0.01 | 0.45 ± 0.04 | 0.70 ± 0.02 | 0.82 ± 0.03 |
|  |  | 2 | 0.41 ± 0.03 | 0.11 ± 0.01 | 0.48 ± 0.04 | 0.74 ± 0.03 | 0.81 ± 0.02 |
|  |  | 5 | 0.46 ± 0.02 | 0.134 ± 0.005 | 0.41 ± 0.02 | 0.70 ± 0.02 | 0.82 ± 0.02 |
| Col-0 |  | 0 | 0.40 ± 0.06 | 0.12 ± 0.02 | 0.48 ± 0.08 | 0.70 ± 0.04 | 0.840 ± 0.009 |
|  |  | 0.25 | 0.42 ± 0.04 | 0.12 ± 0.01 | 0.46 ± 0.05 | 0.68 ± 0.02 | 0.82 ± 0.02 |
|  |  | 1 | 0.47 ± 0.03 | 0.137 ± 0.009 | 0.39 ± 0.04 | 0.70 ± 0.02 | 0.81 ± 0.04 |
|  |  | 2 | 0.46 ± 0.05 | 0.13 ± 0.02 | 0.41 ± 0.07 | 0.74 ± 0.02 | 0.80 ± 0.03 |
|  |  | 5 | 0.45 ± 0.04 | 0.13 ± 0.01 | 0.43 ± 0.05 | 0.72 ± 0.03 | 0.835 ± 0.007 |
| SnRK1.1-*OE* | DCMU | 0 |  |  |  |  |  |
|  |  | 0.25 | 0.04 ± 0.03 | 0.04 ± 0.01 | 0.93 ± 0.04 | 0.2 ± 0.1 | 0.68 ± 0.04 |
|  |  | 1 | 0.02 ± 0.01 | 0.041 ± 0.005 | 0.94 ± 0.01 | 0.09 ± 0.05 | 0.66 ± 0.02 |
|  |  | 2 | 0.032 ± 0.005 | 0.038 ± 0.003 | 0.93 ± 0.01 | 0.17 ± 0.03 | 0.62 ± 0.02 |
|  |  | 5 | 0.04 ± 0.01 | 0.030 ± 0.002 | 0.93 ± 0.01 | 0.31 ± 0.06 | 0.541 ± 0.037 |
| Col-0 |  | 0 |  |  |  |  |  |
|  |  | 0.25 | 0.03 ± 0.04 | 0.04 ± 0.01 | 0.92 ± 0.05 | 0.1 ± 0.2 | 0.68 ± 0.02 |
|  |  | 1 | 0.022 ± 0.006 | 0.038 ± 0.006 | 0.941 ± 0.005 | 0.12 ± 0.05 | 0.62 ± 0.04 |
|  |  | 2 | 0.032 ± 0.005 | 0.037 ± 0.007 | 0.930 ± 0.006 | 0.18 ± 0.06 | 0.62 ± 0.03 |
|  |  | 5 | 0.038 ± 0.005 | 0.034 ± 0.002 | 0.928 ± 0.004 | 0.23 ± 0.03 | 0.56 ± 0.03 |

**Supplementary Table S3.**
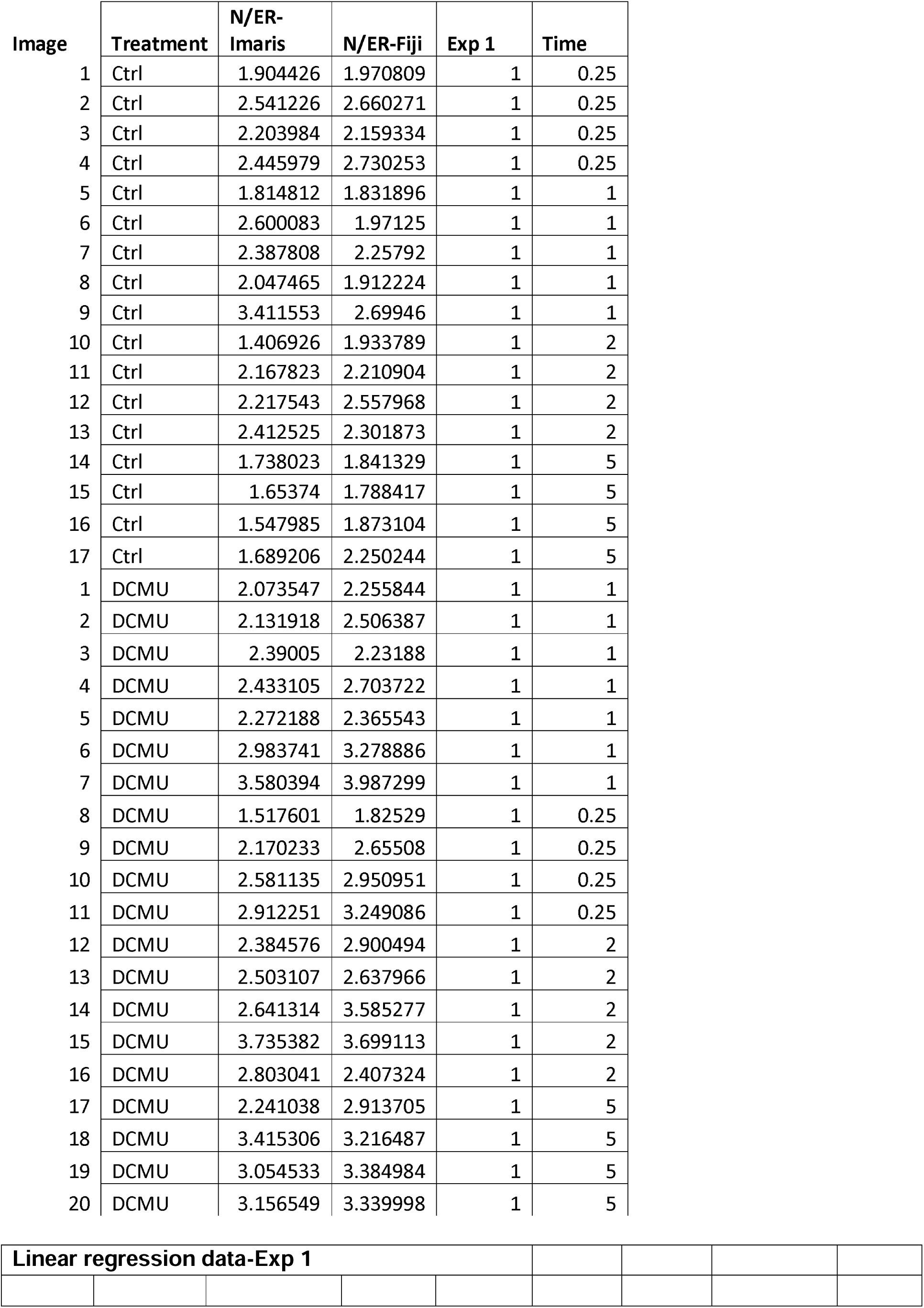
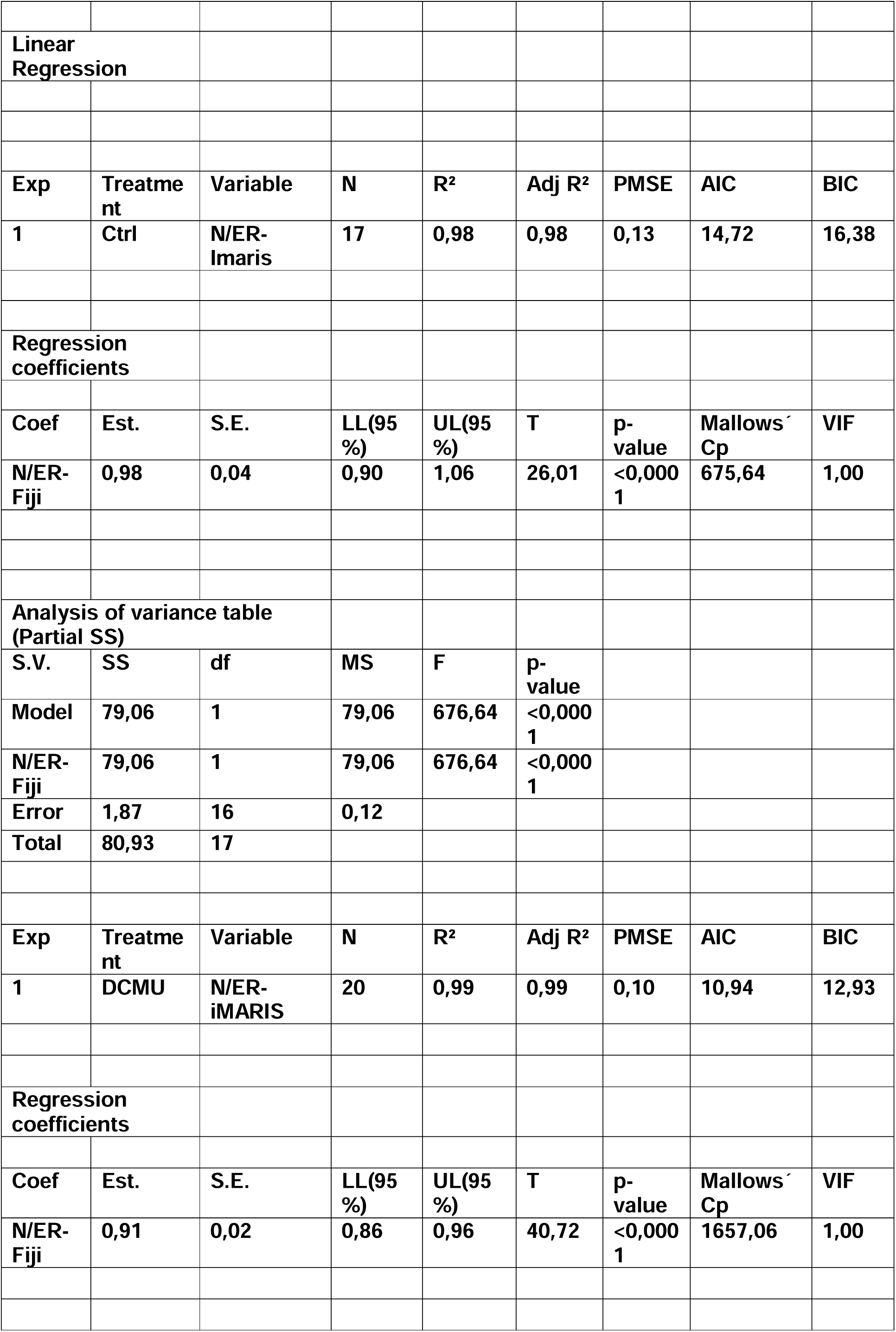
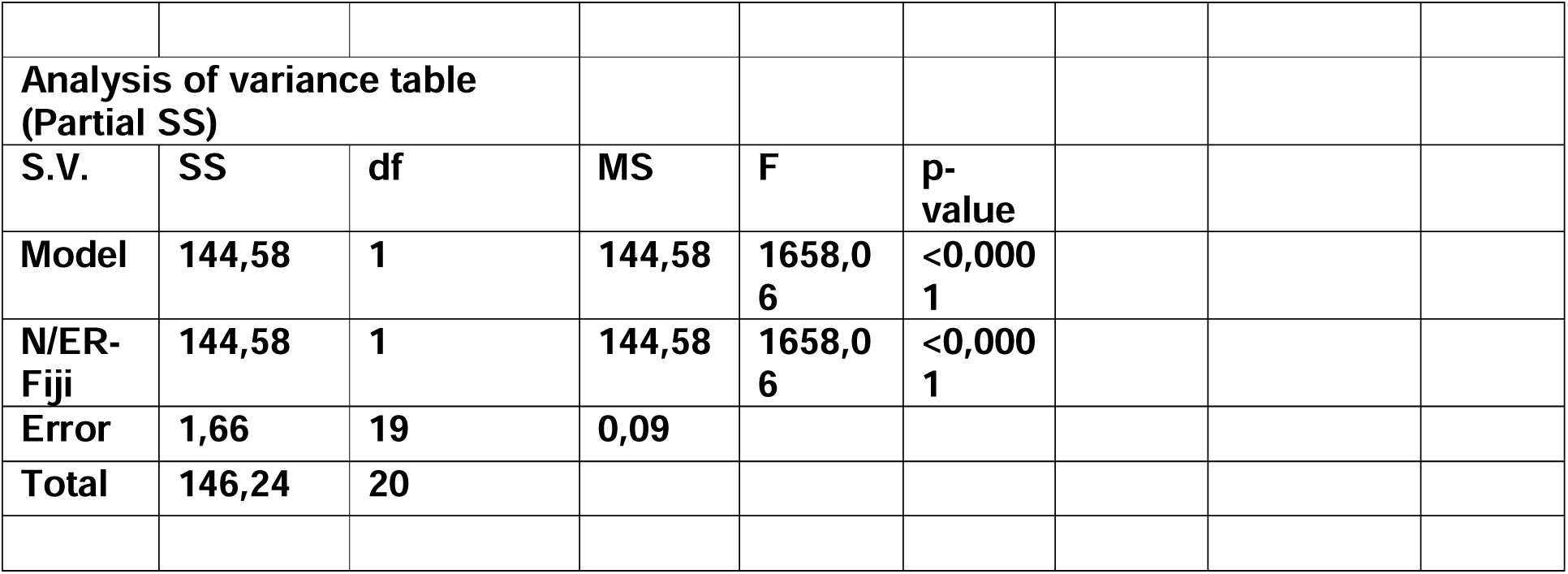
Linear regression.

**Supplementary Figure S1.** Single fluorescence channel. 3D rendering reconstruction of Z-stack images in individual fluorescence channels: SnRK1.1-eGFP (A), ER-RFP (B), and chloroplast auto-fluorescence (C), along with the brightfield image (D). A merged 3D reconstruction combining GFP, ER-RFP, and chloroplast autofluorescence is shown in (E). SnRK1.1 distribution could be analyzed using the SnRK1.1-eGFP channel. ER-RFP channel aid to generate an ER mask for SnRK1.1-ER quantification. Scale bar: 20 μm.

**Supplementary Figure S2.** SnRK1.1-*OE* exhibits a delay in flowering time. (A) Evaluation of flowering time in SnRK1.1 lines: snrk1.1-/- (SnRK1.1 KO lines), Col-0 (wild-type), and SnRK1.1-*OE* lines (SnRK1.1-eGFP fusion lines, #1 and #2). Results show that SnRK1.1-*OE* lines exhibit a significant delay in flowering time (days) compared to Col-0. The *snrk1.1^-/-^* line behaves similarly to wild-type plants. (B) Analysis of the number of rosette leaves revealed no significant differences between SnRK1.1-*OE1*, Col-0, and *snrk1.1^-/-^* lines, except for SnRK1.1-*OE2*, which showed a significant variation. (C) Representative image illustrating differences in flowering time. Sample size (n) = 17. Different letters indicate significant differences (P < 0.05). Scale bar: 10 cm.

**Supplementary Figure S3.** Validation of DCMU effectiveness through photosynthetic measurements. Two-week-old wild-type plants (Col-0) and SnRK1.1-eGFP overexpressing (SnRK1.1-*OE*) plants were treated with DCMU or water (Ctrl). (A) Scheme of the effect of DCMU over the Electron Transport Chain. Electron flow is schemed as a red line. DCMU blocks the electron transport from quinone binding site A (Qa) to site B (Qb) in Photosystem II (PSII). PAM measurements of chlorophyll fluorescence were obtained to calculate (B) fraction of open PSII reaction centers, (C) PSII quantum yield under illumination and (D) maximum quantum yield of PSII in dark-adapted leaves.

**Supplementary Figure S4.** N/ER INDEX experiments resulting from the application of a new protocol. Independent experiments (Experiments 2 and 3) were conducted to quantify SnRK1.1 intracellular distribution under control conditions (Ctrl) and following treatment with 50 μM DCMU over different time points (0.25, 1, 2, and 5 hours). Both graphs illustrate that up to 1 hour post-treatment, the N/ER INDEX increased significantly compared to the control in all time frame.

**Supplementary Figure S5.** OJIP curves for DCMU response. Two-week-old Col-0 and SnRK1.1-*OE* (SnRK1.1-eGFP overexpressors) plants were exposed to 50 μM DCMU (dark line) or Ctrl conditions with water (grey line). The same patterns are observed in both Col-0 (A) and SnRK1.1-*OE* (B) lines under both DCMU and mock treatments. Under DCMU treatment, OJIP curves show a saturated J peak, reflecting the complete reduction of the quinone binding site A in Photosystem II and maximal blockage of downstream electron flow.

